# Aqueous ethanol permeation across Sterlitech flat sheet aquaporin membranes

**DOI:** 10.1101/200709

**Authors:** Jeremy Lewis, Ali Alshami

**Affiliations:** Chemical Engineering – University of North Dakota

## Abstract

Aquaporin (Aqp) embedded membranes have become a topic of recent research due to the superior selectivity of Aqp to water and its acceptable flux. Most researchers have utilized Aqp membranes for water purification purposes such as reverse and forward osmosis desalination of sea and brackish water and wastewater treatment. This paper investigated the permeation of aqueous ethanol across commercial aquaporin membranes for possible applications in ethanol dehydration in the process and biofuel refining industries. The study revealed that rather than being selectively rejected, ethanol was able to diffuse through the membrane with minimal changes in composition from feed to permeate, even at low feed concentrations of ethanol. The total flux across the membranes were shown to average 3.86 liters/m^2^h (LMH). The presence of ethanol increased the relative rate of water transport across the membrane. When comparing total flux of pure water to aqueous ethanol solutions, the flux was higher for solutions containing any ethanol.

## 1. Introduction

### 1.1 Aquaporin (Aqp)

Aqp are a family of transmembrane proteins that are responsible for the selective permeation of water and, depending on the location in the organism, some solutes. Aqp have unique structural features that allow them to be selective and permeable simultaneously. They are helical structures forming an hourglass shape with a narrow constriction site containing histidine and arginine which is responsible for ion rejection [1]. The narrowest point is made up of an asparagine-proline-alanine (NPA) motif which rejects large molecules by steric hindrance and provides a positive electrostatic field that orients and passes water molecules in a single file fashion [2,3].

*In vivo* Aqp exist in an amphiphilic liposome which is part of the cell membrane. The liposome plays an important role in preventing water permeation through the cell membrane itself. *In vitro* it is also desirable and necessary to have Aqp in an amphiphilic carrier to properly orient and stabilize the molecule. Researchers have used liposomes as well as block copolymers as the carrier [3].

### 1.2 AqpZ

Harvesting Aqp from a living organism is no simple task. However, while still non-trivial, the growth and culture of Aqp from E. Coli, termed AqpZ, has shown to be manageable. Hovijitra et al. described a method to synthesize AqpZ into synthetic liposomes [4] and Borgnia described the growth and reconstitution of AqpZ into liposomes, creating “proteoliposomes” or protein incorporated liposomes [5]. Several other researchers have performed similar experiments and have incorporated AqpZ into liposomes as well as synthetic block copolymers [6,7]. Due to this, AqpZ has become the molecule typically used *in vitro*.

### 1.3 Aqp based biomimetic membranes

For many years, researchers have produced membranes that aimed to mimic the cell membrane water selective channels. This class of membrane has been termed “biomimetic” because the Aqp molecule is structurally incorporated into the membrane. Constructing the membrane has also shown to be a non-trivial task. The process followed typically involves dispersing the Aqp into an amphiphilic carrier, either a liposome or block copolymer. Then rupturing the resulting cyclic product onto a substrate, and depositing the substrate onto a porous support [7]. Several methods have been proposed to accomplish this. The most common method is the vesicular rupture method. First, vesicles are absorbed onto a substrate. For this to work properly, the vesicles and substrate should be of opposite charge. As the concentration of vesicles increases, the cyclic vesicles eventually rupture and self-assemble on the substrate in a planar fashion [8]. The substrate is then deposited onto a solid support such as gold [9], alumina [10], or polymer membranes [11]. What these membranes lacked were stability. They could not withstand high pressures for commercial use.

To improve stability many methods have been proposed. One was to add a cushion between the substrate and the porous support. Wang et al. showed that a carboxylated polyethylene glycol polymer cushion improved the flexibility of membranes supported on porous alumina [12]. Another technique that has been used is pore spanning. In this case, proteoliposomes cover the pores of the support material [13]. Another method involved the production of a dense layer over the support [14]. Several other methods have been studied including crosslinking supports [15] and imprinting proteoliposomes into a support [16].

### 1.4 Aqp membranes for water treatment

*In vivo* Aqp are water selective and reject charged species well, so the natural extension was to use this technology for desalination purposes. Researchers have shown salt, specifically sodium chloride and magnesium chloride, are rejected across Aqp based membranes within in the range of 32.9% to 99% and a flux range from 2-34 LMH/bar [1,7]. These studies were typically done on a small scale with membrane surface areas less than 30 cubic centimeters and around 4-5 bar. Commercial Aqp membranes have become available. A company called Aquaporin A/S produced a membrane called Aquaporin Inside^™^. This membrane was made by a hollow fiber module or flat sheet coated by a thin film containing the active Aqp. Another company called Applied Biomimetic produced a membrane with Aqp embedded into a polymer matrix available in a flat sheet and spiral-wound form [7].

### 1.5 Aqueous ethanol separation

When used as a solvent or produced, ethanol is typically in aqueous form. Aqueous ethanol has little market value and cannot be used as a fuel. When ethanol is used as a solvent in a chemical process or is produced from fermentation into a biofuel, the next step is to separate it from solution and recover as much and as pure as possible. However, the separation of ethanol from water is thermodynamically limited to its azeotropic concentration which is about 95% ethanol by mass. To achieve a higher purity ethanol, it is necessary to use techniques beyond simple binary distillation or flash. Typically, the preferred method is absorption or adsorption. Some adsorbents that have proved to be successful are cornmeal [17], lingo-cellulose based materials [18], and zeolite [19]. Although useful, adsorption results in a high pressure drop and columns need to be regenerated periodically. Another method that has been used is distillation with an added entrainer.

Entrainers are able to break the binary ethanol water azeotrope by creating a tertiary azeotrope that that occurs at a lower temperature than the aqueous ethanol [20]. While entrainment distillation is effective, it is energy intensive and some entrainers can be harmful. Another novel method is pervaporation. In pervaporation, the aqueous ethanol is partially vaporized and put into contact with a membrane in which the membrane selectively allows permeation of ethanol [21]. Although less energy intensive than distillation, pervaporation still requires the vaporization of solution. Lastly, there are hybrid methods that combine several other methods into one [22].

### 1.6 Aqp membranes for aqueous ethanol separation

Although Aqp membranes have never been studied, to the best of the authors’ knowledge, for aqueous ethanol separation, some *in vivo* studies have been performed with a variety of results. For example, Lahajnar concluded that ethanol at low concentrations may impede water flux across membrane water channels in bovine red blood cells [23]. Bodis showed that when rat stomachs were subject to ethanol, Aqp levels spiked after five minutes then quickly dropped below initial concentrations and steadily rose slightly higher than initial concentrations [24]. Madeira showed that water flux in yeast membranes across Aqp increased with increasing concentrations of ethanol but became leaky to protons [25]. With these studies it is difficult to determine how ethanol interacts with Aqp and especially does not address Aqp *in vitro*.

It has been speculated, but never proven, that ethanol can transport through Aqp channels [26]. However, looking at the structure, it is apparently intuitive that ethanol should not be able to transport across Aqp. The widest part of the Aqp channel is about 4Å and narrows to approximately 2Å at the selectivity filter [27]. Sterically, this opening is sufficient for water, which is about 2.75Å in diameter. Ethanol is much larger at about 4.4Å in diameter, so its transport should be hindered. However, the transport across Aqp is more intricate than simple pore diffusion. Molecular dynamic simulations such as the one performed by Smolin, suggest the side chains in the Aqp help align molecules so they are electrostatically forced through the channel [28]. Water and ethanol have similar polarities, so it is possible the electrostatic mechanism is more dominant than the steric hindrance.

### 1.7 Research Scope and Objectives

The objective then, of this research is to investigate the transport of aqueous ethanol across commercially made Aqp membranes. It is hypothesized that the transport of ethanol across the membranes will be inhibited. If steric hindrance is the dominant channeling mechanism, the rate of transport of water will be much larger than the rate of transport of ethanol. The permeate would be water rich, ideally pure, and the retentate would be ethanol rich.

## 2. Materials and Methods

### 2.1 Chemicals

190 proof (95% by volume) Everclear grain alcohol was purchased from a local grocery store. The membranes used in these experiments were bought from Sterlitech, as a supplier, and were manufactured by Aquaporin A/S as Aquaporin flat sheets designed for forward osmosis mode. The membranes were rated for at least 7 LMH with a salt water feed solution. The membranes consisted of an Aqp Z-polyamide matrix. Before use, they were stored in a refrigerator between 2-8°C. Each membrane was used only once as to eliminate the possibility of contamination, excessive drying (as per product specifications), and reduced Aqp activity. Concentrated sulfuric acid (95.0-98.0%) and ACS grade potassium dichromate fine crystals were purchased from Fisher Scientific. High purity compressed oxygen gas was supplied by Praxair.

### 2.2 Permeation Apparatus

A low pressure reverse osmosis apparatus was utilized in this study. Membranes were supported by a 47 mm stainless steel filter holder made by Milllipore. The membrane support scheme was adjusted, to preserve Aqp active layer, from manufacturers recommendations as so. The mesh support was positioned at the bottom layer, followed by the membrane with the active side facing up, and a rubber O-ring sealed the gap between the membrane edges and the top fitting. A mesh tube connected the membrane holder to a stainless steel tank used to hold the solution. Oxygen was used to pressurize the fluid in the tank, and the pressure was recorded by a gauge located at the membrane support. The membrane holder was rated for a maximum inlet pressure of 19 bar, and all runs were done at 4 bar.

### 2.3 Qualitative determination of ethanol presence and a variation of the potassium dichromate reagent – preparation

Considering the expected permeate would have little ethanol, a qualitative method was one method used to determine the presence of ethanol. A potassium dichromate reagent as described by Seo, with modification, was used [29]. 8 mL of sulfuric acid was mixed with 20 mL of distilled water then carefully added to 0.85g of potassium dichromate over and icy water bath.

Samples were prepared by pipetting 100 μL of permeate sample. This was mixed with 1 mL of dichromate reagent and diluted with distilled water to a final volume of 10 mL. Any shade of orange indicated low to no concentration of ethanol was present (<20% by mass). Shades of green indicated medium concentration (20-70% by mass), and blue was indicative of high concentration (>70% by mass). Alternatively, a refined reagent could have been used in combination with UV-Vis spectroscopy to have a linearity range in the desired concentration and absorbance measured at a wavelength of 600nm. In these experiments, the reagent was only used at a qualitative measurement to determine if solution concentrations should be quantitatively determined.

### 2.4 Quantitative determination of ethanol concentration

Ethanol concentrations in the permeate were determined quantitatively using FTIR. A zinc selenide crystal was utilized. Distilled water was used as a background sample. Ethanol standards were prepared using the grain alcohol and a series of two distinctive peaks were apparent between wavenumbers 900 and 1100 cm-1. The peak heights were used to calibrate the standards, and were linear over 0-85% by mass, and reported concentrations are a result of the average obtained by comparing both peak heights to calibration.

## 3. Results and Discussion

### 3.1 Permeation Results

Permeation experiments were done at several different feed solution concentrations. Figure 1 shows the feed solution concentration compared to the permeate concentration. The observations show an interesting tend.

**Figure 1:**
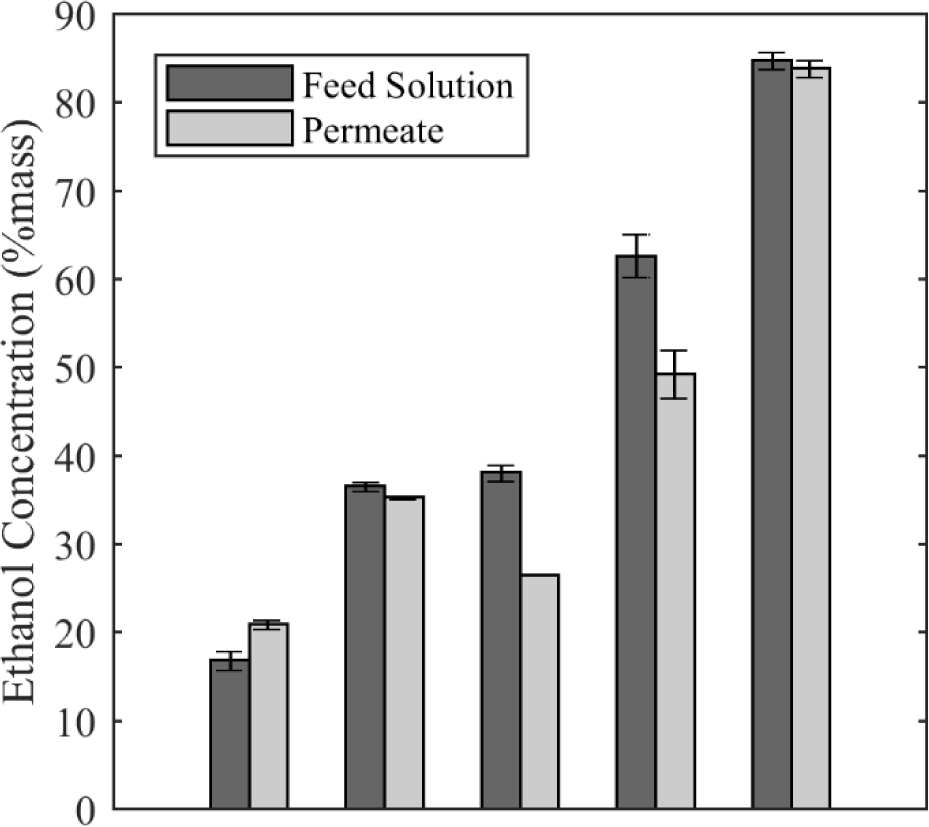
Results from permeation tests. Feed solution was prepared at concentrations between 19 and 80% by mass and the permeate across the membrane was measured. All tests were run at a pressure of 4 bar.

Firstly, separation was not seen in any trial as predicted, this is mostly water permeating the membrane. The rate of ethanol transport appears about equal to the rate of water transport across the membrane. There does not appear to be any practically significant separation. None the less, some insight is gained. At a feed solution of 19% ethanol, the amount of ethanol in the permeate increased slightly. This would indicate the rate of ethanol transport was slight higher than the rate of water transport. Contrarily, at feed solutions between 35-60% the concentration of ethanol in the permeate decreased slightly, indicating the opposite. At 80% ethanol in the feed, the concentrations remained about equal. This would suggest that as the concentration of ethanol in the feed increased, the flux of ethanol decreased compared to water. However, at high concentrations of ethanol, the flux remains similar. From a molecular probability view these results seem counterintuitive. When fewer molecules of ethanol are present in solution, the probability of an ethanol molecule finding an Aqp channel is much less than the probability of water finding a channel. So, the flux of water should be greater at low ethanol concentrations. The opposite was seen. However, these results seem consistent with Madeira, who showed the flux of water across yeast cells increased as ethanol concentrations increased [25].

Another explaination for this phenomena is that mechanism responsible for permeation was across the polyamide matrix, and not the Aqp molecule. From these results, it is apparent that the hypothesized rejection capability of the Aqp molecule was either non existant or out weighed by the lack of rejection provided by the polyamide matrix.

### 3.2 Flux Results

The flux was calculated based on the exposed membrane surface area in the membrane holder and the amount of permeate collected in about one hour. Figure 2 shows the results of these calculations.

**Figure 2:**
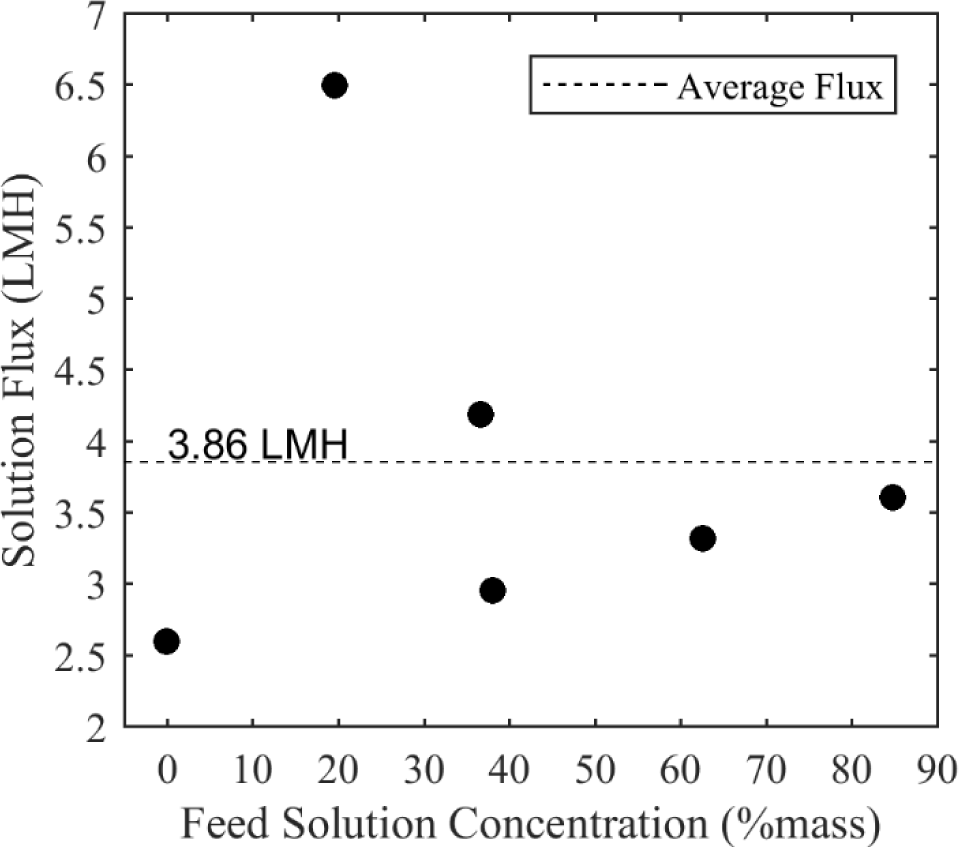
Total solution flux in Liter/m2/h (LMH). Membrane surface area was 13.85 cm2 and the solution was allowed to permeate at 4 bar for about.

It is important to note that the membranes were rated for >7 LMH when exposed to a salt solution in forward osmosis mode. The average taken here of 3.86 LMH was comparatively lower than these specifications. Although, this value is not unreasonable at 4 bar. Xia reported a pure water flux of 0.52 LMH/bar [30], which is comparable to the pure water flux of 0.65 LMH/bar shown here.

It is also important to note that the presence of ethanol increased the total flux slightly, by at least 0.5 LMH. This also supports the *in vivo* results and suggests the transport across Aqp *in vivo* follows a similar mechanism *in vitro* as in the membranes.

### 3.3 Limitations of Work

While the results as presented appear coherent, it is unclear whether the permeation of ethanol occurred across the Aqp channel, the amphiphilic layer, or the polyamide matrix. Literature suggests the ethanol is capable of transporting across polyamide membranes and the polyamide layer can be the limiting factor in rejection [30,31]. Without further experimentation, it cannot be stated definitely, but it is possible the ethanol slipped through the polyamide layer.

Following recommendations from the membrane specifications to soak the membranes for at least thirty minutes before running, after the membranes were run without soaking, they were run again. This would be equivalent to presoaking the membranes for one hour. Doing so had no effect as the resulting flux values were essentially equivalent.

In this study, the membranes used were designed for forward osmosis mode, but were used in a low pressure reverse osmosis cell to reduce the number of variables in the study. Although the pressure used was low, it is possible running the membranes in forward osmosis mode would yield slightly different results.

## 4. Concluding Remarks

### 4.1 Conclusion

The permeation of aqueous ethanol was investigated over commercial Aquaporin flat sheet membranes. The separation of ethanol from water was minimal at all feed concentrations. The rate of water transport across the membranes appeared to increase, and was relatively higher than the rate of transport of ethanol, as the concentration of ethanol increased. The total flux of solution across the membranes increased when comparing pure water flux to the flux of any aqueous ethanol solution.

Most importantly, it is apparent that the steric hindrance hypothesized to be provided by the Aqp channel was not the dominant selectivity mechanism. The electrostatic forces aligning polar molecules and forcing them through the pore, or the transport across the polyamide matrix are possible sources that limited the selectivity.

### 4.2 Future Work

It would be beneficial to investigate alcohols that are more and less polar and larger and smaller in diameter than ethanol. Doing so would provide more insight into the dominant selectivity mechanism. Other aqueous solutions with non-alcoholic polar solutes would also provide more detail. Micrographs should also be taken to determine if, after expose to aqueous ethanol, the surface morphology of the membrane changes. Along with this, the concentration of permeate should be taken at periodic increments to determine if the ethanol and water flux remain constant throughout the experiment or if the ethanol concentrations in the permeate change from start to finish. Lastly, running the experiment in forward osmosis mode with a lower than 4 bar pressure gradient would give more insight into the membrane behavior.

## Acknowledgements

The authors thank Aquaporin A/S for providing the membranes. The work was partially funded by ND EPSCoR. The authors declare no conflicts of interest.

## References

1. Chuyan Tang, et al. “Biomimetic aquaporin membranes coming of age” Desalination 368 (2015) pp.89–105

2. Jiang Yong, MA TongHui. “Imprtance of NPA motifs in the expression and function of water channel aquaporin-1” Chinese Science Bulletin (2007)

3. Yoshinori Fujiyoshi et al. “Structure and Function of Water Channels” Current Opinion in Structural Biology 12.4 (2002) pp. 509–515

4. Normal T. Hovijitra, et al. “Cell-Free Synthesis of Functional Aquaporin Z in Synthetic Liposomes” Biotechnology and Bioengineering 104 (2009) pp. 40–49

5. Mario J. Borgnia, et al. “Functional Reconstitution and Characterization of AqpZ, the E. coli Water Channel Protein” Journal of Molecular Biology 291 (1999) pp.1169–1179

6. Joachim Habel, et al. “Aquaporin-Based Biomimetic Polymeric Membranes: Approaches and Challenges” Membranes 5 (2015) p. 307–351

7. A. Giwa, et al. “Biomimetic membranes: A critical review of recent progress” Desalination 420 (2017) pp.403–424

8. Gregory J. Hardy, Rahul Nayak, Stefan Zauscher. “Model cell membranes: Techniques to for complex biomimetic supported lipid bilayers via vesicle fusion” Current Opinion in Colloid & Interface Science 18.5 (2013) pp. 448–458

9. Anne L. Plant. “Self0Assembled Phospholipid/Alkanethiol Biomimetic Bilayers on Gold” Langmuir 9 (1993) pp. 2764–2767

10. Phuoc H.H. Duong, et al. “Planar biomimetic aquaporin-incorporated triblock copolymer membranes on porous alumina supports for nanofiltration” Journal of Membrane Science 404-410 (2012)pp 34–43

11. Xuesong Li, et al. “Preparation of high performance nanofiltration (NF) membranes incorporated with aquaporin Z” Journal of Membrane Science 450 (2014) pp. 181–188

12. Honglei Wang, et al. “Preparation and characterization of pore-suspending biomimetic membranes embedded with Aquaporin Z on carboxylated polyethylene glycol polymer cushion” Soft Matter 16 (2011) pp. 7274–7280

13. H. Wang, et al. “Highly permeable and selective pore-spanning biomimetic membranes embedded with aquaporin Z.” Small 8.8 (2012) pp. 1185–1190

14. Y. Kaufman, et al. “Towards supported bola amphiphile membranes for water filtration: Roles of lipid and substrate” Journal of Membrane Science 457 (2014) pp. 50–61

15. Hong Lei Wang, et al. “Mechanically robust and highly permeable AquaporinZ biomimetic membranes” Journal of Membrane Science 434 (2013) pp. 130–136

16. Wenyuan Xie, et al. “An aquaporin-based vesicle-embedded polymeric membrane for low energy water filtration” Journal of Materials Chemistry A 26 (2013) pp. 7592–7600

17. Xien Hu. “Fixed-Bed Adsorption and Fluidized-Bed Regeneration for Breaking the Azeotrope of Ethanol and Water” Separation Science and Technology 36.1 (2001) pp. 125–136

18. Tracy J. Benson and Clifford E. George. “Cellulose Based Adsorbent Materials for the Dehydration of Ethanol Using Thermal Swing Adsorption” 11.1 (2005) pp. 697–701

19. Marian Simo, et al. “Adsorption/Desorption of Water and Ethanol on 3A Zeolite in Near Adiabatic Fixed Bed” Industrial & Engineering Chemistry Research 48.20 (2009) pp. 9247–9260

20. R. Katzen, P.W. Madson, G.D. Moon, Jr. “Ethanol distillation: the fundamentals” Katzen International, Inc.

21. H. Eustache, G. Histi. “Separation of aqueous organic mixtures by pervaporation and analysis by mass spectrometry or a coupled gas chromatograph-mass spectrometer” Journal of Membrane Science 8.2 (1981) pp. 105–114

22. Ana M. Eliceche, et al. “Optimization of azeotropic distillation columns combined with pervaporation membranes” Computers and Chemical Engineering 26.4-5 (2002) pp. 563–573

23. Gojmir Lahajnar, Peter Macek, Petra Smid, Ivan Zupancic. “Ethanol-and acetonitrile-induced inhibition of water diffusional permeability across bovine red blood cell membrane” Biochimica et Biophysica Acta 1235 (1995) pp. 437–442

24. Beata Bodis, et al. “Active water selective channels in the stomach: investigation of aquaporins after ethanol and capsaicin treatment in rats” Journal of Physiology-Paris 95 (2001) pp. 271–275

25. Ana Madeira, et al. “Effect of ethanol on fluxes of water and protons across the plasma membrane of Saccharomyces cerevisiae” REMS Yeast Research 10.3 (2010) pp. 252–258

26. Eric Soupene, et al. “Aquaporin Z in Escherichia coli: Reassessment of its Regulation and Physiological Role” Journal of Bacteriology 185.15 (2002) pp. 4304–4307

27. David F. Savage, et al. “Architecture and Selectivity in Aquaporins: 2.5Å X-Ray Structure of Aquaporin Z” PLOS Biology 1.3 (2003)

28. Nikolai Smolin, Bin Li, et al “Side-Chain Dynamics Are Critical for Water Permeation through Aquaporin-1” Biophysical Journal 95.3 (2008) pp. 1089–1098

29. Hyun-Beom Seo, et al. “Measurement of ethanol concentration using solvent extraction and dichromate oxidation and its application to bioethanol production process” Journal of Industrial Microbiology and Biotechnology 36 (2009) pp. 285–292

30. Lingling Xia, et al. “Novel Commercial Aquaporin Flat-Sheet Membrane for Forward Osmosis” Industrial and Engineering Chemistry Research (2017)

31. Meng Shen, et al. “Rejection mechanisms for contaminants in polyamide reverse osmosis membranes” Journal of Membrane Science 509 (2016) pp. 36–47

